# Mountain Centroid: RNA Ensemble Representation with Mountain Profiles

**DOI:** 10.64898/2026.08.19.745640

**Authors:** Takumi Otagaki, Kiyoshi Asai, Kengo Sato

## Abstract

**Background:** RNA molecules form thermodynamic ensembles, but interpretation often requires a single representative structure. Existing base-pair centroid estimators assess agreement at the level of individual base pairs and do not directly target nesting depth along the sequence.

**Methods:** We introduce Mountain Centroid, which minimizes expected squared mountain-profile distance, and derive dynamic programming algorithms with and without RNA pairing constraints. We also combine the Mountain Centroid objective with the base-pair centroid gain.

**Results:** Across 21,254 RNAStrAlign sequences, Mountain Centroid had lower median normalized mean squared mountain distance (NMSMD) than minimum-free-energy (MFE) and base-pair centroid (*γ* = 1) structures, whereas its median base-pair F1 was lower. Imposing RNA pairing constraints improved base-pair F1 for 59.35% of sequences and reduced it for 3.58%. At an illustrative weight, the combined objective had median base-pair F1 similar to MFE while retaining lower median NMSMD than MFE and all tested *γ*-centroid settings.

**Conclusions:** Mountain Centroid represents an RNA structural ensemble with a single secondary structure that reflects how nesting depth varies across nucleotide positions. Combining mountain-profile and individual-base-pair criteria allows their relative contributions to be varied.

## 1. Introduction

RNA structure is closely linked to molecular function and is therefore important for understanding RNA mechanisms (Zhang et al., 2022). Experimental characterization of RNA structure remains challenging, partly because RNA molecules are flexible and can adopt multiple conformations (Deng et al., 2023; Ganser et al., 2019). Computational methods therefore play an important role in predicting RNA structure from sequence. Here, we focus on RNA secondary structure, which describes the pattern of base pairing along the sequence.

RNA secondary-structure analysis often requires a single structure for a given sequence. Under a specified thermodynamic model, an MFE structure minimizes free energy (Zuker and Stiegler, 1981). However, the thermodynamic ensemble includes many possible secondary structures and thus contains information beyond any single MFE structure (McCaskill, 1990). Representing this ensemble with a single structure therefore requires a measure of how well the representative structure reflects the ensemble as a whole, and different measures can emphasize different structural features. Centroid and *γ*-centroid estimators assess this agreement at the level of individual base pairs (Ding et al., 2005; Hamada et al., 2009). However, pair-level agreement does not directly describe how nesting depth varies along the sequence.

The mountain profile records how nesting depth varies across nucleotide positions (Moulton et al., 2000). Because mountain distance compares nesting depth position by position rather than requiring exact base-pair endpoint matches, structures with similar nested architecture can remain close even when some individual base pairs differ. Mountain profiles are therefore useful for comparing nesting patterns across homologous RNAs and identifying regions in which nesting depth is conserved or not. For aligned RNAs, mountain profiles can be compared position by position; this approach has been used to identify evolutionarily conserved structural elements in viral genomes (Hofacker et al., 1998). Ensemble-derived mountain information has also been incorporated into RNA sequence–structure alignment (Bayegan and Clote, 2020). Mountain distance summarizes differences between profiles, providing a measure of overall nesting similarity that complements pair-level agreement. We define Mountain Centroid by assessing ensemble agreement through mountain profiles rather than individual base pairs.

Figure 1 summarizes the mountain representation and illustrates the difference in how MFE and Mountain Centroid structures are defined. An MFE structure has minimum free energy under the thermodynamic model, whereas a Mountain Centroid structure has minimum expected squared mountain distance from the ensemble. These definitions can therefore yield different structures even when they are based on the same thermodynamic model.

**Figure 1.**
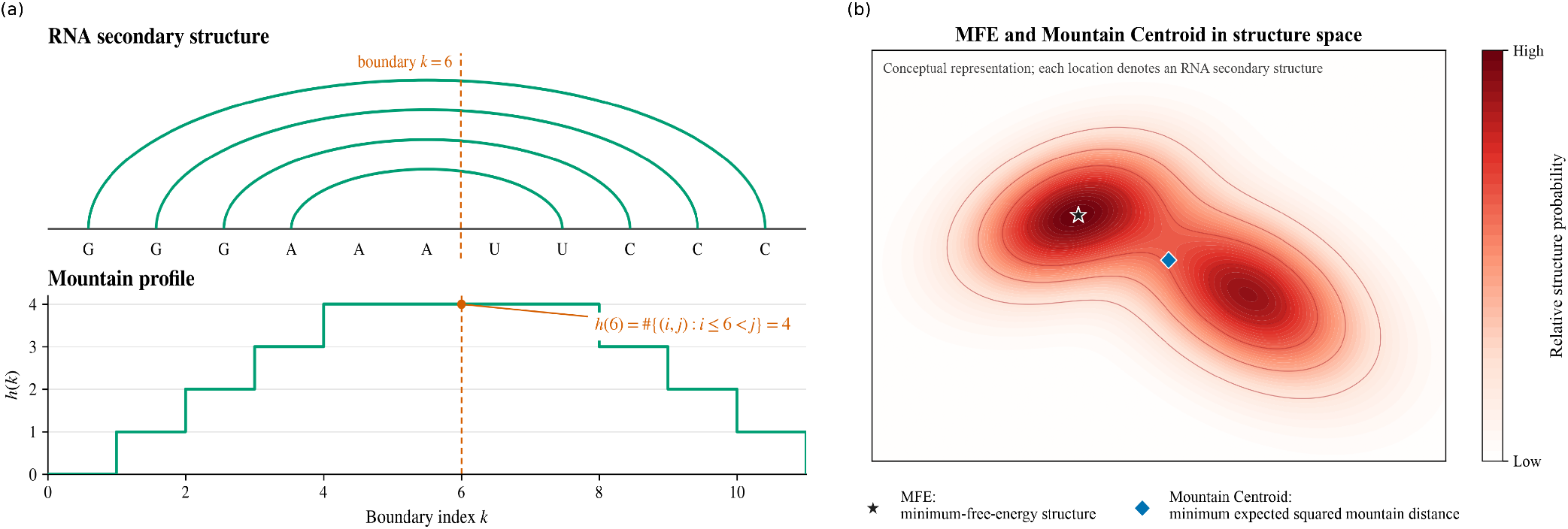
Conceptual overview of Mountain Centroid. (a) An RNA secondary structure and its mountain profile. At the boundary after position *k, h*_*σ*_(*k*) counts the base pairs (*i, j*) satisfying *i* ≤ *k < j*. (b) How MFE and Mountain Centroid structures are defined. An MFE structure has minimum free energy, whereas a Mountain Centroid representative has minimum expected squared mountain distance from the ensemble among candidate structures. The two-dimensional structure space and probability landscape in (b) are schematic and do not represent an explicit embedding.

We develop this representation in two stages. We first formulate a mountain-path relaxation of the objective, and then optimize the same objective over RNA secondary structures satisfying RNA pairing constraints to obtain Mountain Centroid. We then extend it by combining the mountain-profile loss with the base-pair gain used by the centroid estimator (Hamada et al., 2009).

Our contributions are threefold. First, we formulate Mountain Centroid by defining agreement with the ensemble through nesting depth along the sequence. Second, we give algorithms that globally minimize the Mountain Centroid objective over mountain paths and over RNA secondary structures satisfying pairing constraints, and extend the constrained DP to combine mountain-profile and individual-base-pair criteria without changing its time or space complexity. Third, we evaluate Mountain Centroid on the benchmark dataset, use the mountain-path optimization as a comparator to quantify the objective and structural effects of the constraints, and examine how combining the two criteria affects base-pair F1 and NMSMD.

## 2. Methods

### 2.1. Mountain profiles and the ensemble objective

Let *x* = *x*_1_ … *x*_*n*_ be an RNA sequence. We write each base pair as (*i, j*) with *i < j*. Two base pairs (*i, j*) and (*k, ℓ*) cross when *i < k < j < ℓ* or *k < i < ℓ < j*; a structure is pseudoknot-free if it contains no such crossing pair. Throughout, we restrict the ensemble and candidate structures used by the prediction methods to the pseudoknot-free class. Under the nearest-neighbor thermodynamic model, let *σ* be a secondary structure and let *p*(*σ* | *x*) denote its Boltzmann probability for sequence *x* (McCaskill, 1990). Following the mountain representation of RNA secondary structures (Moulton et al., 2000), define the position-indexed profile of a structure *σ* by

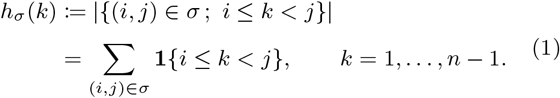

Here, **1**{*C*} denotes the indicator of a condition *C*; it equals one when *C* is true and zero otherwise. Thus *h*_*σ*_(*k*) is the nesting depth immediately after nucleotide position *k*; it counts how many base pairs enclose the boundary between positions *k* and *k* + 1, and hence how deeply that boundary is nested within the structure. A Mountain Centroid structure minimizes the expected squared distance between its mountain profile and the mountain profiles of structures in the ensemble. For structures *σ* and *τ*, define their squared mountain loss as

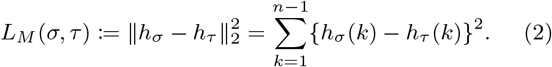

The expected squared mountain loss is minimized by a structure closest to the mean mountain profile.

For a candidate structure or mountain path *σ*, its expected loss over the ensemble is

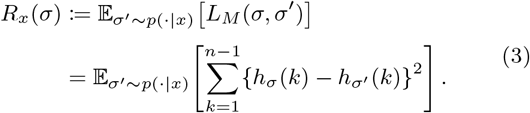

Let 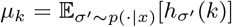 be the expected mountain height at position *k*, and call *µ* = (*µ*_1_, …, *µ*_*n*−1_) the mean mountain profile. For a fixed candidate *σ*, the cross term vanishes because 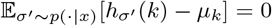, giving

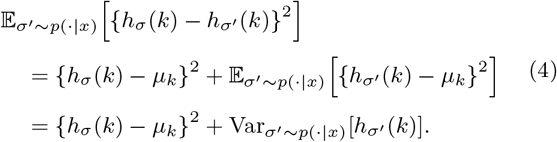

Define the candidate-dependent term

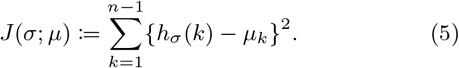

Summing Equation 4 over positions gives

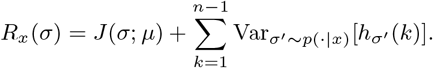

The variance term is determined by the ensemble distribution *p*(· | *x*), not by the candidate structure *σ*. Minimizing the expected loss is therefore equivalent to minimizing *J* (*σ*; *µ*).

We next express *µ* in terms of base-pairing probabilities (BPPs). For *i < j*, define

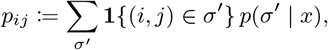

where the sum ranges over all secondary structures *σ*′ in the ensemble. Thus *p*_*ij*_ is the BPP of (*i, j*). Taking expectations in the indicator-sum representation of Equation 1 and using linearity gives

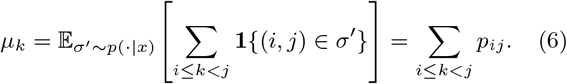

Thus *µ* is obtained directly from the BPPs. This mapping retains the expected nesting depth at each position but discards the identities of the base pairs contributing to that depth. For a pseudoknot-free ensemble under a specified nearest-neighbor thermodynamic model, the McCaskill algorithm, as implemented in ViennaRNA, computes the partition function and BPPs (McCaskill, 1990; Lorenz et al., 2011). The BPP calculation used in the benchmark is specified separately in the Evaluation design.

If nucleotide pairability and the minimum-hairpin requirement are not imposed, the objective in Equation 5 can be minimized over valid nonnegative mountain paths. Let *A*_path_ be the set of base-pair sets encoded by height sequences *h*_0_, …, *h*_*n*_ with *h*_0_ = *h*_*n*_ = 0, *h*_*k*_ ≥ 0, and *h*_*k*_ − *h*_*k*−1_ ∈ {−1, 0, 1}. Steps of +1, 0, and −1 represent 5′ endpoints, unpaired nucleotides, and 3′ endpoints, respectively. An optimal structure over this set satisfies

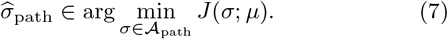

Set *µ*_*n*_ := 0, which does not change *J* because every complete path has *h*_*n*_ = 0. Let *D*[*k, t*] be the minimum accumulated objective over valid partial paths that end at height *t* after position *k*, with *D*[0, 0] = 0 and infeasible states assigned infinite cost. For 1 ≤ *k* ≤ *n* and 0 ≤ *t* ≤ min(*k, n* − *k*), the recurrence is

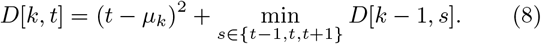

Let *H*_path_ be the maximum number of height levels considered at any position, including height zero. This dynamic program runs in *O*(*nH*_path_) time and uses *O*(*nH*_path_) memory, which are both *O*(*n*^2^) in the worst case. The full state definition, boundary conditions, and traceback procedure are given in Supplementary Text S1 and Algorithm S1.

### 2.2. Mountain Centroid with RNA pairing constraints

For Mountain Centroid, we restrict candidate structures to the set *A*_RNA_ of pseudoknot-free structures in which every base pair (*i, j*) is an AU, UA, GC, CG, GU, or UG pair and satisfies *j* − *i* − 1 ≥ TURN, where TURN = 3. Thus, at least three nucleotides must lie between paired endpoints. Unless otherwise specified, Mountain Centroid refers to the method that enforces the two RNA pairing constraints: nucleotide pairability and the TURN = 3 minimum-hairpin requirement. Let *ℬ*_WC*/*GU_ denote these six ordered Watson–Crick or GU wobble pairs. A Mountain Centroid structure 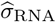 satisfies

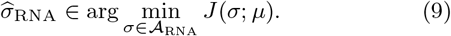

Every candidate in *A*_RNA_ is also represented in *A*_path_, so *A*_RNA_ ⊆ *A*_path_. Consequently, 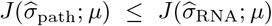. We therefore refer to *σ*_path_ as the mountain-path relaxation of Mountain Centroid. Its objective value is a lower bound on the optimal Mountain Centroid objective value.

The mountain-path recurrence cannot directly enforce the RNA pairing constraints because its state records only the current height, or the number of previous nucleotides that must still be paired with later nucleotides. Determining whether a later nucleotide can pair with one of them would also require their positions and nucleotide identities. Storing this information would create too many states as the sequence grows. Under the pseudoknot-free assumption, crossing pairs are excluded, allowing the sequence to be divided into intervals that can be optimized separately.

For 1 ≤ *k* ≤ *n* and *d* ≥ 0, define the local squared-profile cost at position *k* and height *d* by *c*_*k*_(*d*) := (*d* − *µ*_*k*_)^2^. Define *F* [*i, j, d*] as the minimum possible value of 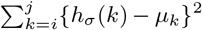 within the interval [*i, j*], given that the interval is enclosed by *d* base pairs whose endpoints lie outside [*i, j*]. We call *d* the external mountain height. Empty intervals have zero cost.

Now consider the leftmost nucleotide *i*. If it is unpaired, the height after position *i* is *d*, so its local cost is *c*_*i*_(*d*) and the remaining interval retains external height *d*. If *i* pairs with a valid position *ℓ*, the pair (*i, ℓ*) spans positions *i*, …, *ℓ* − 1, adding one to the mountain height after each of those positions. Its 5′ endpoint therefore has local cost *c*_*i*_(*d* + 1), the interval [*i* + 1, *ℓ* − 1] has external height *d* + 1, its 3′ endpoint has term *c*_*ℓ*_(*d*), and the interval [*ℓ* + 1, *j*] has external height *d*. These two possibilities give the following recurrence, where *V*_*i*_(*j*) = {*ℓ* : *i* + TURN + 1 ≤ *ℓ* ≤ *j*, (*x*_*i*_, *x*_*ℓ*_) ∈ ℬ_WC*/*GU_} is the set of valid partners of *i* within [*i, j*]: membership in ℬ_WC*/*GU_ specifies the nucleotide-pair type, while the lower bound on *ℓ* requires at least TURN nucleotides between the paired endpoints.

When this set is empty, the inner minimum is interpreted as ∞:

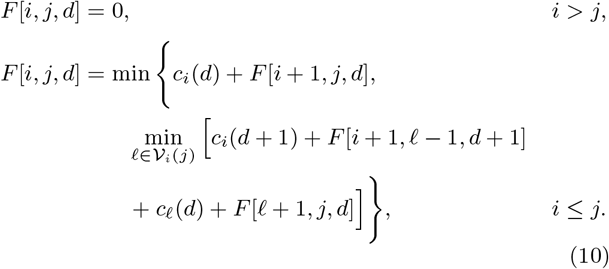

The full objective is *F* [1, *n*, 0], because no pair lies outside the complete sequence. The two recursive intervals in the paired case cannot interact: a pair connecting them would cross (*i, ℓ*). Consequently the recurrence enumerates every valid choice for the leftmost nucleotide without creating a crossing pair.

Correctness follows by induction on interval length. In any nonempty pseudoknot-free interval, its leftmost nucleotide is either unpaired or has one partner *ℓ*. In the latter case, every remaining pair lies wholly inside (*i, ℓ*) or wholly to its right. Equation 10 considers both cases and every valid *ℓ*. Using the best solution for each smaller interval therefore gives the best solution for [*i, j*], completing the induction. The states can be computed top down with memoization, beginning at *F* [1, *n*, 0], so only reachable states are computed. The recursion depth is *O*(*n*), which is dominated by the memoized DP states in the memory bound below. Let *H*_RNA_ be the number of external-height levels reached by this recurrence, including height zero. There are *O*(*n*^2^*H*_RNA_) states, each examining at most *O*(*n*) partners. Time and memory are therefore *O*(*n*^3^*H*_RNA_) and *O*(*n*^2^*H*_RNA_), respectively. Because *H*_RNA_ = *O*(*n*) in the worst case, these bounds become *O*(*n*^4^) time and *O*(*n*^3^) memory. The complexity bounds for both methods are summarized in Table 1. Pseudocode including memoization and traceback is provided in Supplementary Algorithm S2.

**Table 1.** Time and memory complexity. *H*_path_ and *H*_RNA_ denote the numbers of height levels, including zero. The pairing constraints comprise nucleotide pairability and the TURN = 3 minimum-hairpin requirement.

| Method | Pairing constraints | Time | Memory |
| --- | --- | --- | --- |
| Mountain-path relaxation | No | $O(nH_{\text{path}})$ | $O(nH_{\text{path}})$ |
| Mountain Centroid | Yes | $O(n^3 H_{\text{RNA}})$ | $O(n^2 H_{\text{RNA}})$ |

### 2.3. Combining the Mountain Centroid objective with centroid gain

Mountain Centroid evaluates a structure in terms of its mountain-profile distance from the ensemble, whereas the *γ*-centroid estimator evaluates it in terms of the presence or absence of individual base pairs. For a candidate structure *σ* and a structure *σ*′ drawn from *p*(· | *x*), let TP(*σ, σ*′) be the number of base pairs shared by the two structures, and let TN(*σ, σ*) be the number of pairs of nucleotide positions that form a base pair in neither structure. The *γ*-centroid estimator maximizes the expected gain 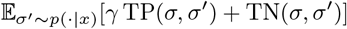 (Hamada et al., 2009).

Using the BPPs, this expected gain can be written as

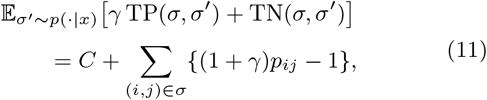

where *C* does not depend on the candidate structure *σ*. Therefore, maximizing the expected gain is equivalent to maximizing the candidate-dependent sum in Equation 11.

We fix *γ* = 1, corresponding to the base-pair centroid estimator. The candidate-dependent centroid gain is therefore

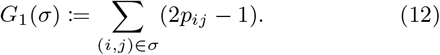

The two criteria have different scales that depend on sequence length. To reduce their dependence on sequence length, we define

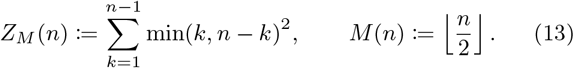

Because both *h*_*σ*_(*k*) and *µ*_*k*_ lie between zero and min(*k, n* − *k*), *J* (*σ*; *µ*)*/Z*_*M*_ (*n*) lies in [0, 1]. A length-*n* secondary structure contains at most *M* (*n*) base pairs, and each term in *G*_1_(*σ*) lies in [−1, 1]; hence *G*_1_(*σ*)*/M* (*n*) lies in [−1, 1].

With *γ* fixed at 1, we use a single parameter *α* ∈ [0, 1] to combine the normalized criteria in the hybrid objective

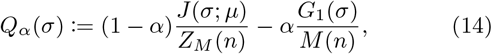

and define

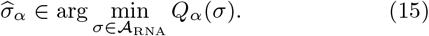

The minus sign converts maximization of the centroid gain into minimization. At *α* = 0, Equation 15 has the same minimizers as Mountain Centroid. At *α* = 1, it gives the base-pair centroid estimator (*γ* = 1) over the same candidate set *A*_RNA_.

The local mountain-profile contribution to *Q*_*α*_ is

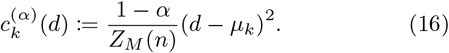

Let *F*_*α*_[*i, j, d*] denote the minimum hybrid-objective contribution of the interval [*i, j*] at external mountain height *d*, analogously to *F* [*i, j, d*] in Equation 10. When the recurrence chooses a base pair (*i, ℓ*), its centroid-gain contribution is added in the corresponding paired branch. The recurrence is

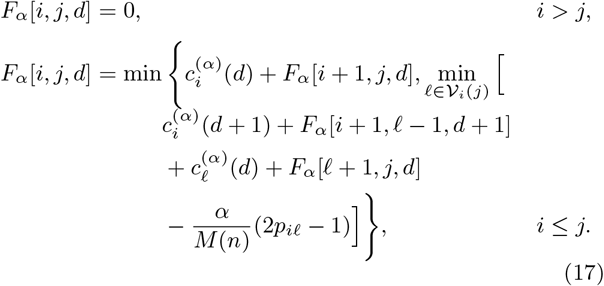

The minimum hybrid objective value is *F*_*α*_[1, *n*, 0]. Equation 17 uses the same interval and external-height variables and the same valid-partner set as Equation 10. It can therefore be evaluated with memoization, and a minimizing structure can be recovered using the same memoization and traceback scheme as in Supplementary Algorithm S2. The numbers of states and partners examined per state are unchanged, so the time and memory complexities remain *O*(*n*^3^*H*_RNA_) and *O*(*n*^2^*H*_RNA_), respectively.

### 2.4. Evaluation design

We evaluated 21,254 RNAStrAlign sequences of at most 300 nt and their reference secondary structures (Tan et al., 2017), as distributed through MultiMolecule (Chen and Zhu, 2024). For each sequence, we computed MFE and centroid structures with the ViennaRNA Package (Lorenz et al., 2011) and a Mountain Centroid structure.

To use ViennaRNA consistently across methods, we obtained BPPs with its implementation of the McCaskill algorithm. We also computed *γ*-centroid structures for *γ* ∈ {0.25, 0.5, 1, 2, 4, 8, 16} using these BPPs; *γ* = 1 corresponds to the centroid estimator. We used these values to examine how the results vary with *γ*. Mountain Centroid and the path relaxation used the same BPP-derived mean profiles. For these mean profiles, both methods find a global optimum over their respective candidate sets. Dataset filtering and calculation settings are described in Supplementary Text S2; software versions and the execution environment are listed in Supplementary Table S1.

For the hybrid objective, we kept *γ* = 1 and evaluated 13 values of *α*. To provide finer resolution near *α* = 0, we specified the values through

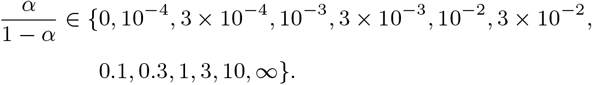

The endpoint odds 0 and ∞ correspond to *α* = 0 and *α* = 1, respectively. We did not use agreement with the reference structures to select a default *α*.

LinearPartition provides an optional faster BPP calculation, approximating BPPs in *O*(*nb*^2^) time for beam size *b* (Zhang et al., 2020). When *b* is fixed, the *O*(*n*^2^) mountain-path DP sets the overall worst-case time of the path-relaxation calculation. The Mountain Centroid DP retains its *O*(*n*^4^) worst-case bound regardless of how the BPPs are computed.

The Mountain Centroid objective *J* in Equation 5 measures squared distance to the ensemble mean profile. The direct objective cost of imposing the RNA pairing constraints is

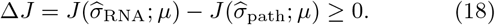

The inequality follows because *A*_RNA_ ⊆ *A*_path_. Structural agreement with the reference secondary structure *σ*_ref_ was assessed using base-pair F1 and normalized mean squared mountain distance (NMSMD). Writing *P*_*σ*_ for the set of base pairs in *σ*, we defined

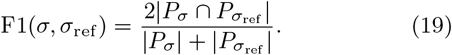

A pair counted as a match only when both endpoints matched the reference pair. To assess agreement between mountain profiles, we used

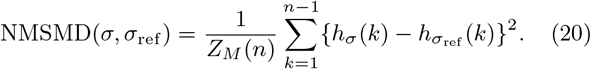

Here *Z*_*M*_ (*n*) is the length-dependent normalizer defined in Equation 13, so NMSMD lies in [0, 1]. This normalization does not change the ordering of results for a fixed sequence. The two quantities use different target profiles. The objective *J* in Equation 5 measures squared distance from the ensemble mean profile, whereas NMSMD in Equation 20 measures normalized squared distance from the reference profile. Thus, the RNA pairing constraints cannot decrease *J*, but they may either increase or decrease NMSMD. All method comparisons are paired by sequence.

## 3. Results

### 3.1. Mountain and base-pair agreement favor different methods

Across all 21,254 benchmark records, including homologous and duplicate sequences, median NMSMD relative to the reference secondary structures was 0.00568 for Mountain Centroid, compared with 0.00647 for MFE and 0.00734 for centroid (*γ* = 1) (Figure 2; Supplementary Table S2). The path relaxation had a median NMSMD of 0.00565. Median base-pair F1 was 0.553 for Mountain Centroid, 0.712 for MFE, 0.716 for the base-pair centroid (*γ* = 1), and 0.500 for the path relaxation. Of the seven tested *γ* values, *γ* = 2 gave the lowest median NMSMD (0.00693) and the highest median base-pair F1 (0.718) for *γ*-centroid structures (Figure 2c). Mountain Centroid had a strictly lower NMSMD than MFE for 67.34% of sequences; the corresponding fraction for the path relaxation was 67.39%. These results show that matching the reference mountain profile and recovering its individual base pairs are related but different goals. The lower NMSMD should therefore be interpreted as closer agreement between mountain profiles, not as an overall improvement in secondary-structure prediction or exact base-pair recovery. In the deduplicated sensitivity analysis (16,222 sequences), Mountain Centroid and the path relaxation had lower median NMSMD than MFE and centroid (*γ* = 1), whereas MFE and centroid (*γ* = 1) had higher median base-pair F1. In six of the seven families, both Mountain Centroid and the path relaxation had lower median NMSMD than MFE and centroid; either MFE or centroid had the highest median base-pair F1 in every family (Supplementary Table S2).

**Figure 2.**
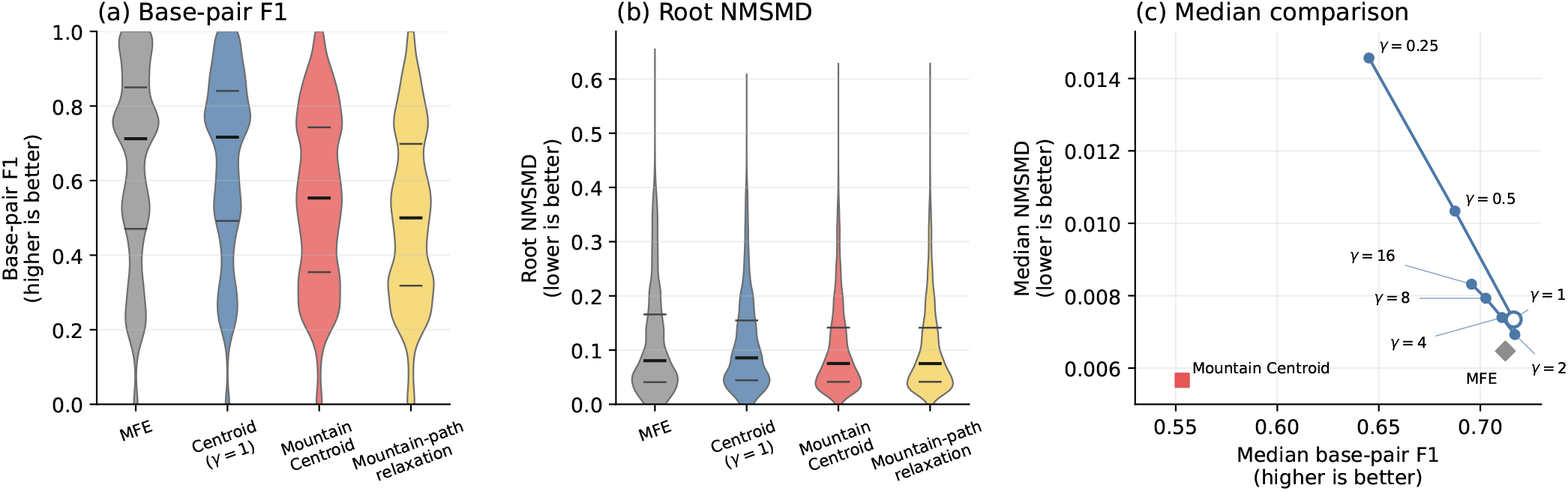
Agreement with reference secondary structures in the full RNAStrAlign benchmark (21,254 records, including homologous and duplicate sequences). (a,b) Distributions of base-pair F1 and root NMSMD 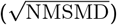. Wider sections of each violin indicate a higher relative concentration of observations at that value; thick black horizontal lines show medians, and thinner horizontal lines show the first and third quartiles. Mountain Centroid, MFE, and centroid (*γ* = 1) are RNA structure predictions. The path relaxation is a lower-bound comparator for *J* and may violate nucleotide pairability or the minimum-hairpin requirement. (c) Median base-pair F1 and median NMSMD for *γ*-centroid structures at the seven tested *γ* values, with MFE and Mountain Centroid shown for comparison. Each point summarizes the same 21,254 records, and the line connects *γ* settings in increasing order.

For an illustrative 5S rRNA sequence, Mountain Centroid had lower NMSMD than both MFE and centroid (0.00316 versus 0.00449), but its base-pair F1 was lower (0.635 versus 0.769; Supplementary Text S6 and Figure S6).

Only 2,767 of 21,254 path-relaxation results (13.0%) satisfied both RNA pairing constraints. Imposing the constraints improved base-pair F1 for 59.35% of sequences, left it unchanged for 37.08%, and reduced it for 3.58%. The median increase in the Mountain Centroid objective, normalized by the maximum possible squared mountain distance for the sequence length, was 1.98 *×* 10^−5^ (Supplementary Text S5 and Figure S5).

Excluding ViennaRNA BPP computation, the Mountain Centroid calculation took a median of 0.0142 seconds per sequence and a maximum of 1.334 seconds during the 15-worker TSUBAME4.0 run. These times include program startup and result parsing.

We also examined the BPP assigned to every base pair present in each reference or predicted structure. Considering all base pairs together, Mountain Centroid structures had a median BPP of 0.914, but 17.64% of their base pairs had BPP below 0.01; the corresponding values for reference structures were 0.834 and 8.99% (Supplementary Text S3 and Figure S1). Supplementary Table S3 reports the median fraction of base pairs with BPP below 0.01 separately for each RNA family.

### 3.2. Base-pair F1 and NMSMD vary with the hybrid weight

Across the 13 tested *α* values, increasing the contribution of centroid gain increased median base-pair F1 from 0.553 at *α* = 0 to 0.716 at *α* = 1, while median NMSMD generally increased from 0.00568 to 0.00734 (Figure 3). At *α* = 0.0909, median base-pair F1 was 0.7077 and median NMSMD was 0.005940. At *α* = 0.2308, the corresponding values were 0.7143 and 0.006098. At *α* = 0.2308, median base-pair F1 was close to that of MFE (0.7124) and the base-pair centroid with *γ* = 2 (0.7179), while median NMSMD remained lower than for MFE (0.006472) and every tested *γ*-centroid setting (the lowest was 0.006927 at *γ* = 2). For 14,704 of 21,254 sequences (69.2%), base-pair F1 was at least as high as for MFE and NMSMD was no greater than for MFE. This value of *α* is presented as an illustrative point on the curve, not as a selected default.

**Figure 3.**
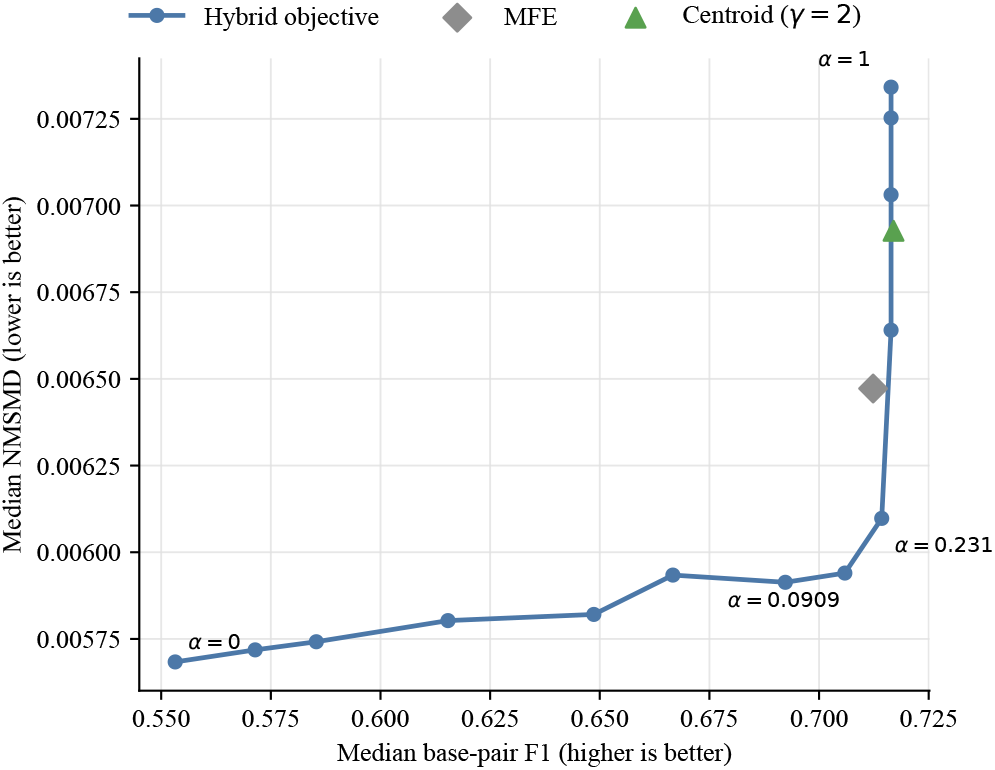
Effect of combining mountain-profile loss and centroid gain in the full RNAStrAlign benchmark (21,254 records). The blue curve shows median base-pair F1 and median NMSMD at the 13 tested *α* values, connected in increasing order of *α*. The endpoint *α* = 0 corresponds to Mountain Centroid, whereas *α* = 1 corresponds to the base-pair centroid (*γ* = 1). MFE and the base-pair centroid (*γ* = 2) are shown as comparison points. Labels identify the two endpoints and two illustrative interior values.

At *α* ≈ 0.001 and 0.003, all seven RNA families had higher median base-pair F1 than at *α* = 0 in both the full and deduplicated analyses. Changes in median NMSMD differed among families (Supplementary Figure S2 and Table S4).

Across the hybrid curve, the fraction of records whose structure contained at least one predicted pair with BPP below 0.01 decreased from 79.21% at *α* = 0 to 1.35% at *α* = 0.0909 and 0.0094% (2 of 21,254 records) at *α* = 0.2308 (Supplementary Text S4 and Figure S3).

At the same *α*, 68.63% of records had hybrid base-pair F1 at least as high as and NMSMD no greater than the corresponding values for the base-pair centroid at *γ* = 2. The hybrid result was better on at least one metric and worse on neither for 43.22% of records. The base-pair centroid at *γ* = 2 was better on at least one metric and worse on neither for 11.04% of records (Supplementary Text S4 and Figure S4).

## 4. Discussion

The choice of an ensemble representative depends on which structural features are considered most important. The base-pair centroid estimator uses agreement on individual base pairs, whereas Mountain Centroid uses the nesting-depth pattern across nucleotide positions. The MFE structure provides a different reference point by minimizing free energy under the thermodynamic model. Varying *γ* affected both median base-pair F1 and median NMSMD, but none of the tested *γ*-centroid settings had a median NMSMD as low as Mountain Centroid. Because the mean mountain profile does not retain base-pair identities, Mountain Centroid can include pairs with low individual BPPs. This behavior was observed in the benchmark, although low-BPP pairs also occurred in reference structures.

The hybrid objective combines mountain-profile loss with base-pair centroid gain without imposing a hard BPP threshold. Small positive values of *α* reduced low-BPP pairs and improved median base-pair F1 across all seven RNA families, with only small changes in median NMSMD. Because there is no generally applicable rule for choosing *α*, the results are interpreted as showing how the two metrics vary across the tested weights rather than as defining a default predictor.

Imposing the RNA pairing constraints increased the Mountain Centroid objective only slightly after normalization, while improving or preserving base-pair F1 for most sequences (Supplementary Text S5 and Figure S5).

Because ViennaRNA was used to compute the MFE and centroid structures and the BPPs for Mountain Centroid, differences among the outputs reflect the definitions of the representatives rather than differences in upstream software. Applying the *O*(*n*^4^) Mountain Centroid algorithm to substantially longer RNAs will require further optimization.

Mountain Centroid provides a mountain-profile-based representative of an RNA structural ensemble and may support future analyses of mountain profiles and ensemble-level nesting patterns. Its hybrid extension combines mountain-profile loss and base-pair centroid gain within the same DP framework.

## Acknowledgements

This study was carried out using the TSUBAME4.0 supercomputer at Institute of Science Tokyo. OpenAI Codex was used for language editing, consistency checks, LaTeX and figure preparation, and code review. The authors reviewed all outputs and take full responsibility for the manuscript and analyses.

## Funding

This work was supported by the Japan Society for the Promotion of Science (JSPS) KAKENHI Grant Numbers 26KJ1158 to T.O., JP24H00737 to K.A. and K.S., and 25H01166 to K.S.

## Data and software availability

Benchmark code and summary results are available at github.com/TakumiOtagaki/MountainCentroidBenchmark. The reusable Mountain Centroid implementation is available at github.com/TakumiOtagaki/MountainCentroid.

## Notes

https://github.com/TakumiOtagaki/MountainCentroid

https://github.com/TakumiOtagaki/MountainCentroidBenchmark

### Competing Interest Statement

The authors have declared no competing interest.

## References

A. H. Bayegan and P. Clote. RNAmountAlign: Efficient software for local, global, semiglobal pairwise and multiple RNA sequence/structure alignment. PLoS One, 15(1):e0227177, Jan. 2020.

Z. Chen and S. Y. Zhu. MultiMolecule, 2024.

J. Deng, X. Fang, L. Huang, S. Li, L. Xu, K. Ye, J. Zhang, K. Zhang, and Q. C. Zhang. RNA structure determination: From 2D to 3D. Fundam. Res., 3(5):727–737, Sept. 2023.

Y. Ding, C. Y. Chan, and C. E. Lawrence. RNA secondary structure prediction by centroids in a boltzmann weighted ensemble. RNA, 11(8):1157–1166, Aug. 2005.

L. R. Ganser, M. L. Kelly, D. Herschlag, and H. M. Al-Hashimi. The roles of structural dynamics in the cellular functions of RNAs. Nat. Rev. Mol. Cell Biol., 20(8):474–489, Aug. 2019.

M. Hamada, H. Kiryu, K. Sato, T. Mituyama, and K. Asai. Prediction of RNA secondary structure using generalized centroid estimators. Bioinformatics, 25(4):465–473, Feb. 2009.

I. L. Hofacker, M. Fekete, C. Flamm, M. A. Huynen, S. Rauscher, P. E. Stolorz, and P. F. Stadler. Automatic detection of conserved RNA structure elements in complete RNA virus genomes. Nucleic Acids Res., 26(16):3825–3836, Aug. 1998.

R. Lorenz, S. H. Bernhart, C. Höner Zu Siederdissen, H. Tafer, C. Flamm, P. F. Stadler, and I. L. Hofacker. ViennaRNA package 2.0. Algorithms Mol. Biol., 6(1):26, Nov. 2011.

J. S. McCaskill. The equilibrium partition function and base pair binding probabilities for RNA secondary structure. Biopolymers, 29(6-7):1105–1119, May 1990.

V. Moulton, M. Zuker, M. Steel, R. Pointon, and D. Penny. Metrics on RNA secondary structures. J. Comput. Biol., 7 (1-2):277–292, Feb. 2000.

Z. Tan, Y. Fu, G. Sharma, and D. H. Mathews. TurboFold II: RNA structural alignment and secondary structure prediction informed by multiple homologs. Nucleic Acids Res., 45(20): 11570–11581, Nov. 2017.

H. Zhang, L. Zhang, D. H. Mathews, and L. Huang. LinearPartition: linear-time approximation of RNA folding partition function and base-pairing probabilities. Bioinformatics, 36(Suppl 1):i258–i267, July 2020.

J. Zhang, Y. Fei, L. Sun, and Q. C. Zhang. Advances and opportunities in RNA structure experimental determination and computational modeling. Nat. Methods, 19(10):1193–1207, Oct. 2022.

M. Zuker and P. Stiegler. Optimal computer folding of large RNA sequences using thermodynamics and auxiliary information. Nucleic Acids Res., 9(1):133–148, Jan. 1981.

