## Supplementary Information for "Mountain Centroid: RNA Ensemble Representation with Mountain Profiles"

### Text S1. Mountain-path relaxation

Let $\mathcal{A}_{\text{path}}$ be the set of base-pair sets encoded by height sequences $h_0, \dots, h_n$ with $h_0 = h_n = 0$, $h_k \geq 0$, and $h_k - h_{k-1} \in \{-1, 0, 1\}$. At each nucleotide, these three possible changes correspond to the $5'$ endpoint of a base pair, an unpaired nucleotide, and the $3'$ endpoint of a base pair, respectively. Under the pseudoknot-free assumption, a stack-based traceback pairs each $3'$ endpoint with the most recent unmatched $5'$ endpoint, yielding a noncrossing base-pair set. The set $\mathcal{A}_{\text{path}}$ does not impose nucleotide pairability or the TURN = 3 minimum-hairpin requirement. Its optimum is the mountain-path relaxation of Mountain Centroid:

$$\widehat{\sigma}_{\text{path}} \in \arg\min_{\sigma \in \mathcal{A}_{\text{path}}} J(\sigma; \mu). \tag{1}$$

Set $\mu_n \coloneqq 0$. Because every candidate satisfies $h_n = 0$, the Mountain Centroid objective can be extended to $k = n$ without changing its value. For $0 \leq k \leq n$ and each integer $0 \leq t \leq \min(k, n-k)$, let $\mathcal{P}_{k,t}$ be the set of partial height sequences $\boldsymbol{h} = (h_0, \dots, h_k)$ that start at $h_0 = 0$, end at $h_k = t$, never fall below zero, and change by at most one at each position. Define

$$D[k, t] \coloneqq \min_{\boldsymbol{h} \in \mathcal{P}_{k,t}} \sum_{r=1}^{k} (h_r - \mu_r)^2. \tag{2}$$

At $k = n$, $\mathcal{P}_{n,0}$ contains all complete mountain paths, so $D[n, 0] = J(\widehat{\sigma}_{\text{path}}; \mu)$. The bound $t \leq n - k$ keeps only partial height sequences that can return to zero by position $n$. Assigning infinite cost to states outside the stated range gives

$$\begin{aligned}
&D[k,t] = 0, \qquad k = 0,\ t = 0,\\
&D[k,t] = \infty, \qquad t < 0 \text{ or } t > \min(k, n-k),\\
&D[k,t] = (t - \mu_k)^2 + \min_{s \in \{t-1, t, t+1\}} D[k-1, s],\\
&\qquad 1 \leq k \leq n, \quad 0 \leq t \leq \min(k, n-k).
\end{aligned} \tag{3}$$

Let $H_{\text{path}}$ be the maximum number of height levels considered at any position, including height zero. The dynamic program has $O(nH_{\text{path}})$ states and requires $O(nH_{\text{path}})$ time and memory. Because $H_{\text{path}} = O(n)$ in the worst case, both are $O(n^2)$. Recording which previous height gives the minimum at each state allows the minimizing path to be reconstructed as detailed in Algorithm S1.

### Algorithm S1. Mountain-path relaxation

The main text defines the mountain-path recurrence using the states $D[k,t]$ and expected profile heights $\mu_k$. These states can be computed bottom up as follows.

---

**Algorithm S1** Mountain-path relaxation

---

**Require:** Expected profile heights $\mu_1, \dots, \mu_{n-1}$
**Ensure:** Globally optimal mountain path and objective value
Set $\mu_n \leftarrow 0$ and $D[0,0] \leftarrow 0$; treat states outside the allowed range as $\infty$
**for** $k = 1$ to $n$ **do**
    **for** $t = 0$ to $\min(k, n-k)$ **do**
        $A \leftarrow \{s \in \{t-1, t, t+1\} : D[k-1, s] < \infty\}$
        $s^\star \leftarrow \arg\min_{s \in A} D[k-1, s]$, breaking ties by the smallest $s$
        $D[k,t] \leftarrow D[k-1, s^\star] + (t - \mu_k)^2$
        $\mathrm{back}[k,t] \leftarrow s^\star$
    **end for**
**end for**
Set $h_n \leftarrow 0$ and trace back from $(n, 0)$ to recover $h_{n-1}, \dots, h_0$
Interpret steps $+1, 0, -1$ as $5'$ endpoints, unpaired nucleotides, and $3'$ endpoints
Pair each $3'$ endpoint with the most recent unmatched $5'$ endpoint
**return** The base-pair set and $D[n, 0]$

---

### Algorithm S2. Mountain Centroid with RNA pairing constraints

The main text defines the constrained recurrence using the states $F[i, j, d]$, local costs $c_i(d)$, and valid-partner sets $\mathcal{V}_i(j)$. These states can be computed top down with memoization as follows.

---

**Algorithm S2** Mountain Centroid with RNA pairing constraints

---

**Require:** Sequence $x_1 \ldots x_n$ and expected heights $\mu_1, \ldots, \mu_{n-1}$, with TURN = 3
**Ensure:** A Mountain Centroid structure and its objective value

Set $\mu_n \leftarrow 0$
**function** Solve$(i, j, d)$ with memoization
**if** $i > j$ **then return** 0
**end if**
**if** $(i, j, d)$ is memoized **then return** $F[i, j, d]$
**end if**
$F[i, j, d] \leftarrow c_i(d) + \text{Solve}(i + 1, j, d)$
back$[i, j, d] \leftarrow$ UNPAIRED
**for** each $\ell \in \mathcal{V}_i(j)$, in increasing order **do**
    $v \leftarrow c_i(d + 1) + \text{Solve}(i + 1, \ell - 1, d + 1) + c_\ell(d) + \text{Solve}(\ell + 1, j, d)$
    **if** $v < F[i, j, d]$ **then**
        $F[i, j, d] \leftarrow v$
        back$[i, j, d] \leftarrow$ PAIR_WITH$(\ell)$
    **end if**
**end for**
Memoize $F[i, j, d]$
**return** $F[i, j, d]$
**end function**
Evaluate Solve$(1, n, 0)$
Trace back from $(1, n, 0)$
**return** The base-pair set and $F[1, n, 0]$

---

### Text S2. Benchmark calculation settings

The RNAStrAlign input contained 37,149 records. After converting T to U, we applied the filters in the order used by the evaluation code: 4,011 records containing characters other than A, C, G, and U were excluded, followed by 11,884 records longer than 300 nt. Retained records were also required to have matching sequence and reference lengths and reference secondary structures containing only `.`, `(`, `)`, `[`, and `]`. No additional records failed these latter requirements. The resulting benchmark contained 21,254 records from seven RNA families. Of these, 97 had pseudoknotted reference structures in which additional base pairs were denoted by square brackets. We preserved these brackets and parsed round and square brackets independently. Base pairs were compared by their endpoints for base-pair F1, irrespective of bracket type. For NMSMD, the reference profile height after each nucleotide position was calculated as the number of reference base pairs spanning that position.

MFE and centroid structures were computed using ViennaRNA Package 2.7.2. Calculations used the package defaults at 37 °C, including dangle model 2, GU pairs, lonely pairs, and special-hairpin corrections. $\gamma$-centroid structures were computed from ViennaRNA BPPs using an independent dynamic-programming implementation. At $\gamma = 1$, its output matched the ViennaRNA centroid for all 21,254 records.

Mountain-path relaxation results and Mountain Centroid structures were computed using Mountain Centroid v1.0.0. Both used the same expected mountain profiles obtained from ViennaRNA BPPs at 37 °C with the same default energy model and dangle settings used for MFE and centroid prediction. The Mountain Centroid solver permitted AU, UA, GC, CG, GU, and UG pairs and used TURN = 3.

Both solvers used deterministic tie-breaking when solutions had equal objective values. When multiple heights at position $k - 1$ gave the same cost for a path-relaxation state at position $k$, the solver selected the smaller height. At the final return to height zero, it therefore selected $h_{n-1} = 0$ rather than $h_{n-1} = 1$ when their costs were equal. For Mountain Centroid, the solver preferred leaving the leftmost nucleotide of an interval unpaired; among tied pairing choices, it selected the partner with the smallest position index.

The illustrative 5S rRNA example was selected deterministically. Eligible records were 80–200 nt, had results from all four calculations, satisfied path-relaxation mean squared mountain distance (MSMD) < Mountain Centroid MSMD < MFE MSMD and path-relaxation F1 < Mountain Centroid F1 < MFE F1, and had Mountain Centroid F1 at least as high as its overall median (0.553). Among the 850 eligible records, we selected the record whose Mountain Centroid–MFE MSMD difference was closest to the median difference among eligible records, with the identifier used to break any tie.

### Table S1. Software and execution environment

Table S1: Software and execution environment.

| Item | Setting |
|---|---|
| Mountain Centroid | v1.0.0 |
| Python | 3.13.13 |
| uv | 0.11.11 |
| NumPy | 2.5.1 |
| ViennaRNA Package | 2.7.2 |
| Matplotlib | 3.11.0 |
| GCC/G++ | 11.5.0 |
| R | 4.4.0 |
| R4RNA | 2.0.9 |
| Compiler settings | C++17 with `-O3 -DNDEBUG` |
| Compute system | TSUBAME4.0 |
| Allocation | 32-CPU interactive environment |
| Mountain Centroid calculation | 15 single-threaded workers |
| Thread control | OpenBLAS, OpenMP, and MKL thread counts fixed to one |
| Output handling | Per-sequence outputs merged after all 21,254 sequences had completed |

### Table S2. Benchmark sensitivity summaries

For the deduplicated sensitivity analysis, records were grouped by RNA sequence. One record was retained when all records for that sequence had the same reference secondary structure; sequences with conflicting references were excluded. This yielded 16,222 unique sequences. Across the full benchmark, the absolute change in NMSMD after imposing the RNA pairing constraints had a median of $9.26 \times 10^{-5}$, an interquartile range of $1.93 \times 10^{-5}$–$2.89 \times 10^{-4}$, and a maximum of 0.00950.

The effects of the RNA pairing constraints were similar after deduplication. Among the 16,222 retained sequences, base-pair F1 improved in 60.72%, was unchanged in 35.75%, and decreased in 3.53%. The Spearman correlation of $\Delta J_{\mathrm{norm}}$ with the F1 change was $\rho = 0.352$, and that of the path-relaxation violation count with the F1 change was $\rho = 0.454$, compared with 0.352 and 0.472, respectively, in the full benchmark.

Table S2: Benchmark results for the full and deduplicated datasets and individual RNA families. Panel A compares medians in the full and deduplicated datasets. Panel B reports medians for each RNA family in the deduplicated dataset; each method cell gives median base-pair F1 followed by median NMSMD.

**A. Full benchmark and deduplicated medians**

| | Full benchmark ($n = 21{,}254$) | | Deduplicated ($n = 16{,}222$) | |
|---|---|---|---|---|
| Method | BP F1 | NMSMD | BP F1 | NMSMD |
| MFE | 0.712 | 0.00647 | 0.696 | 0.00719 |
| Centroid ($\gamma = 1$) | 0.716 | 0.00734 | 0.703 | 0.00782 |
| Centroid ($\gamma = 2$) | 0.718 | 0.00693 | 0.703 | 0.00734 |
| Mountain Centroid | 0.553 | 0.00568 | 0.540 | 0.00607 |
| Mountain-path relaxation | 0.500 | 0.00565 | 0.486 | 0.00599 |

**B. Family-stratified deduplicated medians (BP F1/NMSMD)**

| Family | $n$ | MFE | Centroid ($\gamma = 1$) | Centroid ($\gamma = 2$) | Mountain Centroid | Mountain-path relaxation |
|---|---|---|---|---|---|---|
| 16S rRNA | 142 | 0.655/0.00921 | 0.681/0.00731 | 0.676/0.00815 | 0.460/0.00519 | 0.417/0.00514 |
| 5S rRNA | 9,273 | 0.676/0.00725 | 0.688/0.00738 | 0.696/0.00692 | 0.500/0.00491 | 0.435/0.00493 |
| RNase P | 31 | 0.577/0.00889 | 0.625/0.00786 | 0.600/0.00830 | 0.531/0.00666 | 0.468/0.00669 |
| SRP | 293 | 0.609/0.00768 | 0.641/0.00877 | 0.632/0.00803 | 0.534/0.00694 | 0.485/0.00686 |
| Group I intron | 47 | 0.667/0.00479 | 0.658/0.00479 | 0.672/0.00507 | 0.542/0.00367 | 0.497/0.00367 |
| tRNA | 6,428 | 0.727/0.00705 | 0.723/0.00836 | 0.711/0.00794 | 0.564/0.00796 | 0.533/0.00789 |
| tmRNA | 8 | 0.482/0.01811 | 0.540/0.01601 | 0.508/0.01903 | 0.480/0.01469 | 0.467/0.01474 |

### Text S3. Base-pair probabilities in benchmark structures

To examine the probabilities assigned to individual base pairs, we recorded, for each benchmark record, the base-pair probability (BPP) $p_{ij}$ of every pair $(i, j)$ present in one of five structures: the RNAStrAlign reference secondary structure (Reference), MFE, Centroid ($\gamma = 1$), Centroid ($\gamma = 2$), or Mountain Centroid. BPPs were calculated with ViennaRNA using the settings described in Text S2. The analysis retained all 21,254 records, including homologous and duplicate sequences.

When each base pair was treated as one observation, the median BPP was 0.914 for Mountain Centroid structures, although 17.64% of their pairs had BPP below 0.01. For reference structures, the corresponding values were 0.834 and 8.99%. All Mountain Centroid pairs with BPP below 0.01 were AU, GC, or GU pairs. In the reference structures, 1.54% of pairs below 0.01 were other nucleotide combinations; after excluding them, 8.86% of reference AU, GC, or GU pairs still had BPP below 0.01. The median BPP was 0.887 for MFE structures, and 0.003% of their pairs had BPP below 0.01. These low-BPP pairs occurred in only two MFE structures.

For the $\gamma$-centroid estimator used here, a base pair $(i, j)$ can be included only when

$$p_{ij} > \frac{1}{1 + \gamma}.$$

Consequently, Centroid ($\gamma = 1$) and Centroid ($\gamma = 2$) cannot contain pairs with BPP below 0.5 and 1/3, respectively. In contrast, Mountain Centroid does not impose a pairwise BPP threshold.

To determine whether pairs with BPP below 0.01 were concentrated in only a few records, we calculated their fraction separately for each record. For Mountain Centroid, 79.21% of records contained at least one such pair, and the median fraction across records was 13.6%. For reference structures, 42.27% of records with at least one base pair contained such a pair, and the median fraction was 0. The median per-record fraction for Mountain Centroid was positive in all seven RNA families, ranging from 10.0% to 23.7%, whereas the family medians for MFE and both centroid estimators were 0 (Table S3).

### Figure S1. Base-pair probabilities in benchmark structures

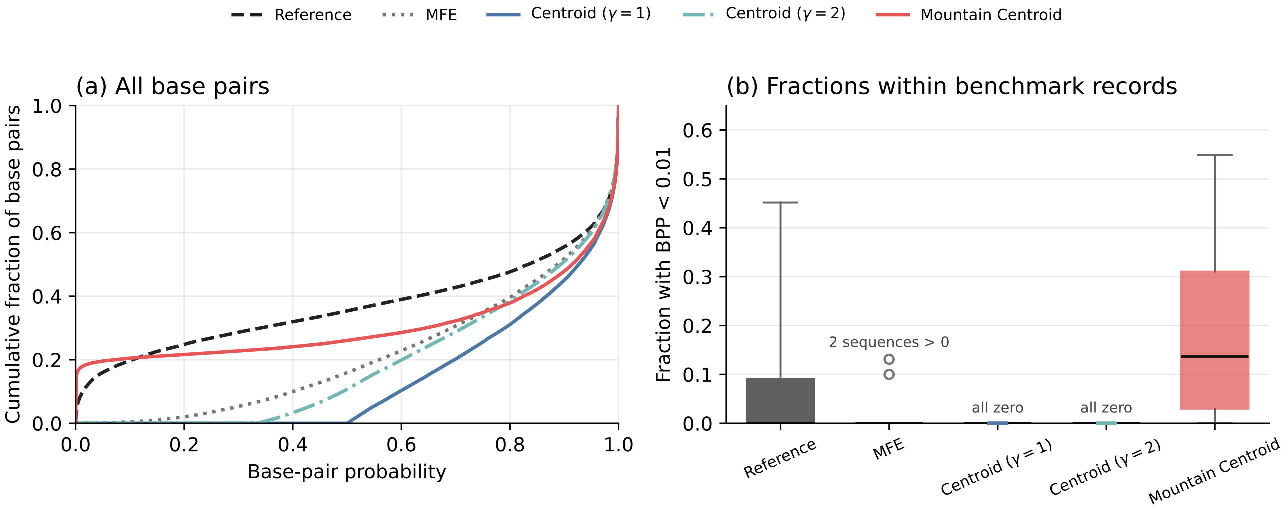


Figure S1: Base-pair probabilities within benchmark structures. BPPs were calculated with ViennaRNA using the settings described in Text S2. Reference denotes the RNAStrAlign reference secondary structure. (a) Across all base pairs in the benchmark, each curve shows the fraction with BPP at or below a given value on the horizontal axis. (b) For each structure containing at least one base pair, we calculated the fraction of its base pairs with BPP below 0.01. Boxes show the middle 50% of these fractions, black lines show medians, and vertical lines extend from the 5th to the 95th percentile. Observations outside the vertical lines are omitted, except for the two nonzero MFE observations shown explicitly. Four reference structures and seven Centroid ($\gamma = 1$) structures contained no base pairs and were omitted from panel (b). Homologous and duplicate sequences were retained.

### Table S3. Low-BPP base pairs by RNA family

Table S3: Median fraction, across records in each RNA family, of base pairs with BPP below 0.01. The $n$ column gives the total number of benchmark records in each family; structures without base pairs were omitted from the relevant median. Homologous and duplicate sequences were retained.

| Family | $n$ | Reference | MFE | Centroid ($\gamma = 1$) | Centroid ($\gamma = 2$) | Mountain Centroid |
|---|---|---|---|---|---|---|
| 16S rRNA | 182 | 11.4% | 0% | 0% | 0% | 23.7% |
| 5S rRNA | 11,302 | 2.9% | 0% | 0% | 0% | 14.3% |
| RNase P | 36 | 8.8% | 0% | 0% | 0% | 20.5% |
| SRP | 407 | 12.9% | 0% | 0% | 0% | 10.0% |
| Group I intron | 83 | 9.5% | 0% | 0% | 0% | 18.6% |
| tRNA | 9,234 | 0% | 0% | 0% | 0% | 12.5% |
| tmRNA | 10 | 11.8% | 0% | 0% | 0% | 16.4% |

#### Text S4. Hybrid objective analysis

We evaluated the hybrid objective for all 21,254 benchmark records at the 13 weights specified in the main text. The calculation used the same BPPs and RNA pairing constraints as Mountain Centroid. Each record was evaluated at every weight, yielding 276,302 structures. At $\alpha = 0$ and $\alpha = 1$, the resulting structures matched Mountain Centroid and the base-pair centroid ($\gamma = 1$), respectively, for all records.

We also summarized the results for the 16,222-sequence deduplicated dataset described in Table S2. Table S4 reports median base-pair F1 and median NMSMD at every tested weight for the full and deduplicated datasets. Figure S2 compares the corresponding curves for all families together and separately for each RNA family. No value of $\alpha$ was selected from agreement with the reference structures as a default.

The changes from $\alpha = 0$ at $\alpha \approx 0.001$ and 0.003 were similar in the full and deduplicated datasets. Relative to $\alpha = 0$ for the same record, the median change in base-pair F1 was +0.0246 at $\alpha \approx 0.001$ and +0.0328 at $\alpha \approx 0.003$ in the full benchmark. The corresponding changes in the deduplicated dataset were +0.0263 and +0.0339. Median changes in NMSMD were 0 and $2.25 \times 10^{-5}$ in the full benchmark and 0 and $2.50 \times 10^{-5}$ after deduplication, respectively. At both weights, median base-pair F1 was higher than at $\alpha = 0$ in all seven RNA families, whereas the changes in median NMSMD differed among families (Figure S2). Thus, the increase in base-pair F1 was not explained solely by repeated identical sequences.

For each $\alpha$, we calculated both the fraction of records whose predicted structure contained at least one base pair with BPP below 0.01 and the median, across records, of the within-structure fraction of such pairs. Structures without predicted base pairs were assigned a value of zero when calculating the median. The fraction of records containing at least one low-BPP pair decreased from 79.21% at $\alpha = 0$ to 1.35% at $\alpha = 0.0909$ and 0.0094% (2 of 21,254 records) at $\alpha = 0.2308$; it was zero at every tested $\alpha \geq 0.5$. The median within-structure fraction decreased from 13.6% at $\alpha = 0$ to 2.7% at $\alpha \approx 0.001$ and was zero by $\alpha \approx 0.003$ (Figure S3).

We also compared the hybrid result with the base-pair centroid at $\gamma = 2$ for each benchmark record at every $\alpha$. At $\alpha = 0.2308$, hybrid base-pair F1 was no lower and hybrid NMSMD was no higher than the corresponding $\gamma = 2$ centroid values for 68.63% of records. This 68.63% consisted of 43.22% for which at least one metric was better for the hybrid result and 25.41% for which both metrics were equal. The $\gamma = 2$ centroid was no worse on either metric and better on at least one for 11.04% of records. For the remaining 20.33%, one method had higher base-pair F1 and the other had lower NMSMD (Figure S4).

### Figure S2. Hybrid objective results before and after deduplication

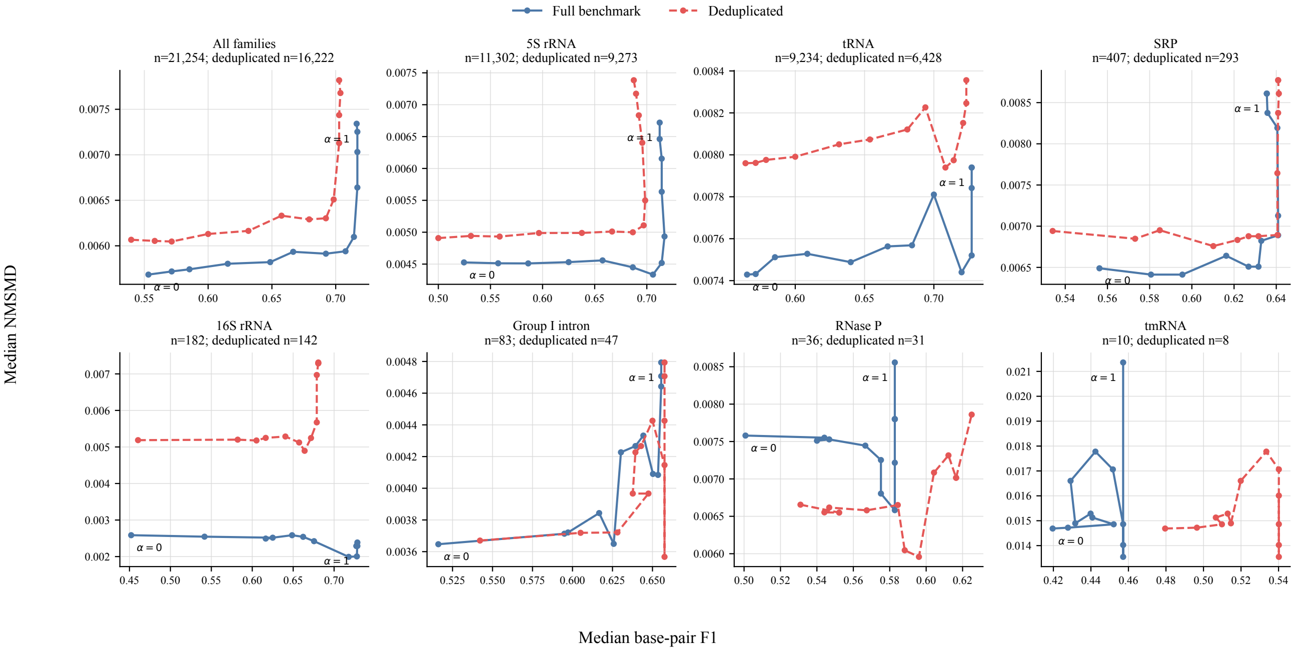


Figure S2: Hybrid-objective results before and after benchmark deduplication. Each panel shows median NMSMD against median base-pair F1 across the 13 tested $\alpha$ values. Solid blue curves show the full benchmark, and dashed red curves show the deduplicated dataset. The solid blue curve in the All families panel is the same 21,254-record curve shown in main-text Figure 3; the dashed red curve adds the corresponding result for the 16,222-sequence deduplicated dataset. The remaining seven panels show the full and deduplicated results separately for each RNA family. Panel titles give the numbers of records in the two datasets. Labels mark $\alpha = 0$ and $\alpha = 1$ on the full-benchmark curves. Higher base-pair F1 is to the right, and lower NMSMD is toward the bottom.

### Table S4. Hybrid objective results across tested weights

Table S4: Median base-pair F1 and NMSMD for the hybrid objective at the 13 tested weights. The first column gives the odds used to specify $\alpha$. The full benchmark contains 21,254 records, including homologous and duplicate sequences. The deduplicated dataset contains the 16,222 sequences described in Table S2. Values of $\alpha$ are rounded to six decimal places.

| | | Full benchmark | | Deduplicated | |
|---|---|---|---|---|---|
| $\alpha/(1-\alpha)$ | $\alpha$ | BP F1 | NMSMD | BP F1 | NMSMD |
| 0 | 0 | 0.5532 | 0.005684 | 0.5397 | 0.006066 |
| $10^{-4}$ | 0.000100 | 0.5714 | 0.005718 | 0.5581 | 0.006054 |
| $3 \times 10^{-4}$ | 0.000300 | 0.5854 | 0.005742 | 0.5714 | 0.006047 |
| $10^{-3}$ | 0.000999 | 0.6154 | 0.005803 | 0.6000 | 0.006130 |
| $3 \times 10^{-3}$ | 0.002991 | 0.6486 | 0.005821 | 0.6316 | 0.006164 |
| $10^{-2}$ | 0.009901 | 0.6667 | 0.005934 | 0.6571 | 0.006331 |
| $3 \times 10^{-2}$ | 0.029126 | 0.6923 | 0.005913 | 0.6792 | 0.006290 |
| 0.1 | 0.090909 | 0.7077 | 0.005940 | 0.6923 | 0.006303 |
| 0.3 | 0.230769 | 0.7143 | 0.006098 | 0.6984 | 0.006509 |
| 1 | 0.500000 | 0.7170 | 0.006640 | 0.7027 | 0.007129 |
| 3 | 0.750000 | 0.7170 | 0.007031 | 0.7027 | 0.007436 |
| 10 | 0.909091 | 0.7170 | 0.007253 | 0.7037 | 0.007679 |
| $\infty$ | 1.000000 | 0.7164 | 0.007341 | 0.7027 | 0.007820 |

#### Figure S3. Low-BPP pairs across hybrid weights

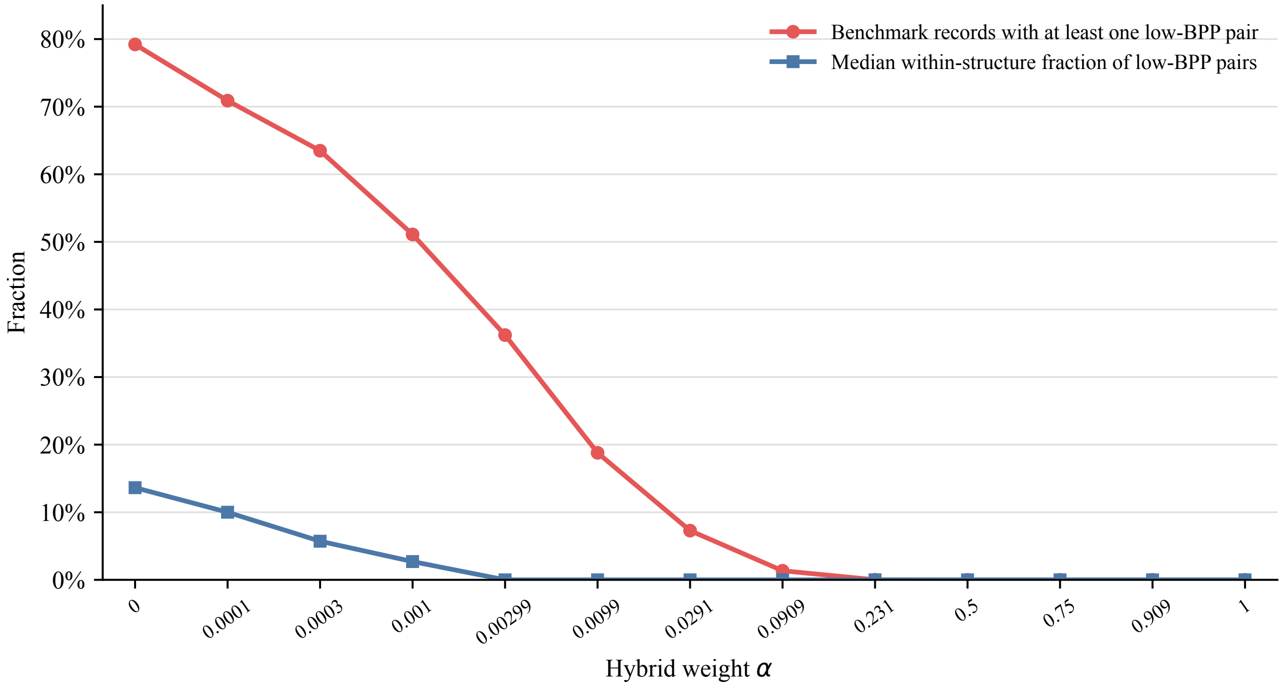


Figure S3: Low-BPP pairs in structures obtained with the hybrid objective across the 13 tested $\alpha$ values. A low-BPP pair was defined as a predicted base pair with BPP below 0.01. Red circles show the fraction of the 21,254 benchmark records whose structure contained at least one low-BPP pair. Blue squares show the median, across records, of the within-structure fraction of predicted pairs with BPP below 0.01; structures without predicted base pairs were assigned a value of zero for this summary. At $\alpha = 0$, the hybrid objective corresponds to Mountain Centroid, whereas $\alpha = 1$ corresponds to the base-pair centroid with $\gamma = 1$. Homologous and duplicate sequences were retained.

### Figure S4. Record-level comparison with base-pair centroid ($\gamma = 2$)

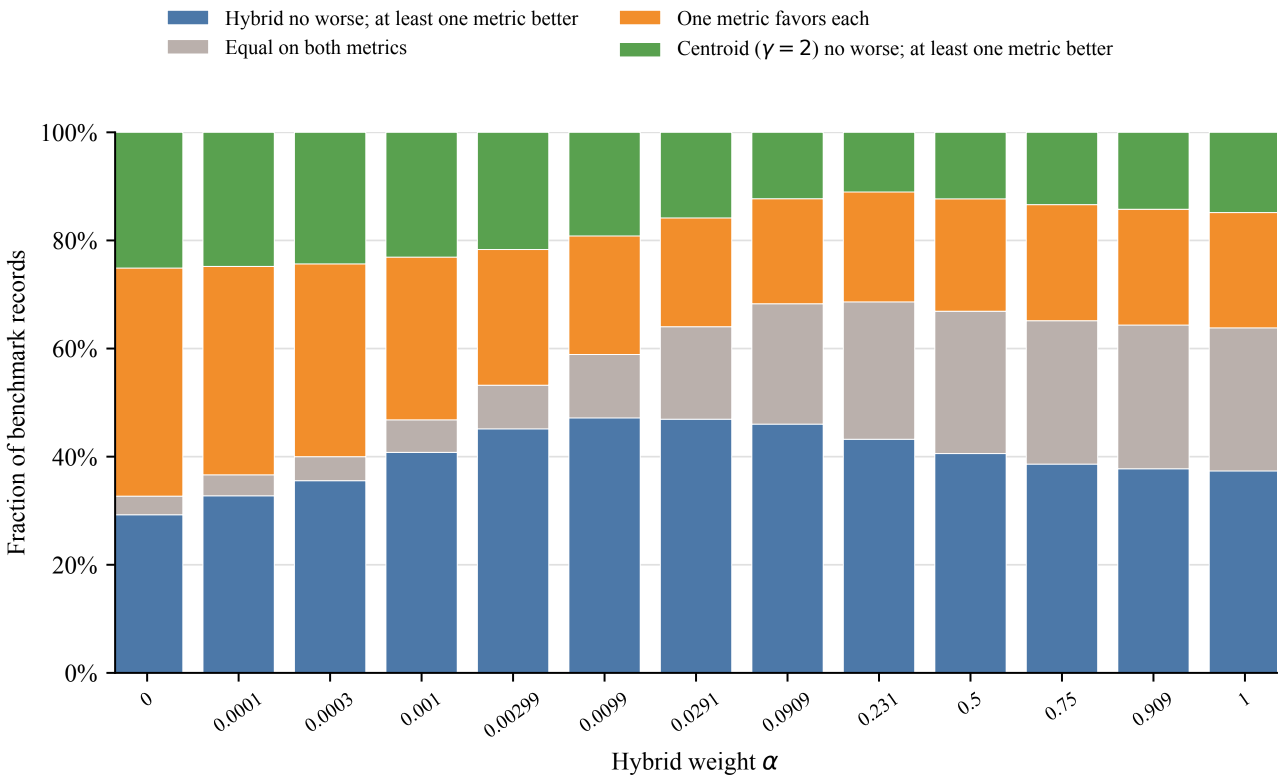


Figure S4: Record-level comparison of the hybrid result with the base-pair centroid at $\gamma = 2$ across the 13 tested $\alpha$ values. Each bar divides the 21,254 benchmark records into four mutually exclusive groups based on base-pair F1 and NMSMD. Blue indicates records for which the hybrid result was no worse on either metric and better on at least one. Green indicates the corresponding condition in favor of the $\gamma = 2$ centroid. Gray indicates equality on both metrics. Orange indicates that one method had higher base-pair F1 and the other had lower NMSMD. Homologous and duplicate sequences were retained.

### Text S5. Effect of RNA pairing constraints

The mountain-path relaxation minimizes the Mountain Centroid objective over a broader candidate set that does not impose nucleotide pairability or the TURN = 3 minimum-hairpin requirement. Only 2,767 of 21,254 path-relaxation results (13.0%) satisfied both constraints. Let $\Delta J = J(\widehat{\sigma}_{\mathrm{RNA}}; \mu) - J(\widehat{\sigma}_{\mathrm{path}}; \mu)$ be the objective increase after imposing the constraints. To compare this increase across sequence lengths, we defined

$$\Delta J_{\mathrm{norm}} = \frac{\Delta J}{\sum_{k=1}^{n-1} \min(k, n-k)^2}. \tag{4}$$

Imposing the constraints increased the objective by a median $\Delta J$ of 1.699 (interquartile range 0.280–4.321). The median $\Delta J_{\mathrm{norm}}$ was $1.98 \times 10^{-5}$ (interquartile range $3.10 \times 10^{-6}$–$5.48 \times 10^{-5}$). The Spearman correlation between $\Delta J_{\mathrm{norm}}$ and the change in base-pair F1 after imposing the constraints was $\rho = 0.352$.

Base-pair F1 improved for 59.35% of sequences, remained unchanged for 37.08%, and decreased for 3.58%. The path relaxation contained a median of five pairs that violated at least one constraint (interquartile range 2–8), and the violation count had a Spearman correlation of $\rho = 0.472$ with the change in F1. When the relaxation already satisfied both constraints, F1 was unchanged in all 2,767 cases. The fraction with improved F1 rose from 27.2% for one violating pair to 57.5% for two or three, 70.8% for four to seven, and 84.4% for eight or more. The median absolute change in NMSMD after imposing the constraints was $9.26 \times 10^{-5}$. The corresponding analysis after benchmark deduplication is summarized in Table S2.

#### Figure S5. Effect of imposing RNA pairing constraints

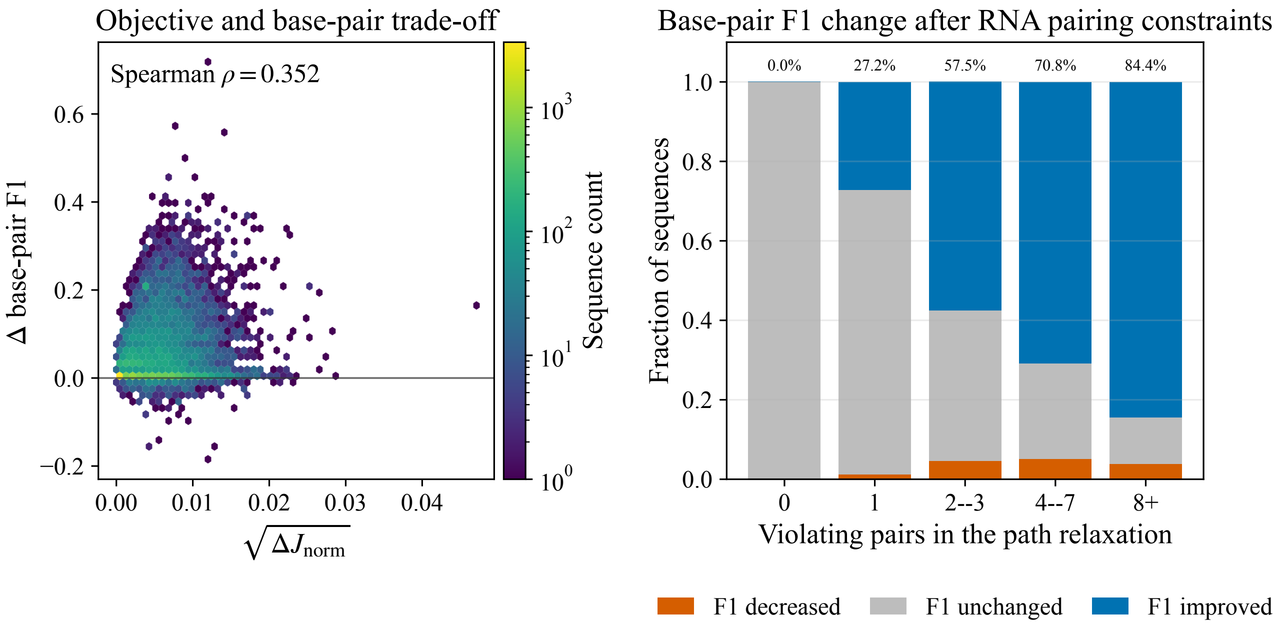


Figure S5: Effect of imposing RNA pairing constraints on 21,254 benchmark records. (a) Square root of the normalized objective increase $\sqrt{\Delta J_{\text{norm}}}$ versus the change in base-pair F1 after imposing the constraints. Hexagon color shows the number of records. (b) Fractions of records in which base-pair F1 decreased, remained unchanged, or improved, grouped by the number of path-relaxation pairs that violated at least one RNA pairing constraint. Percentages above the bars give the fraction with improved base-pair F1. Homologous and duplicate sequences were retained.

#### Text S6. Illustrative 5S rRNA example

The 115-nt bacterial 5S rRNA sequence `5S_rRNA-Bacteria-B00665` illustrates how mountain-profile and base-pair agreement can favor different structures (Figure S6). Imposing the RNA pairing constraints increased NMSMD relative to the reference secondary structure from 0.00297 to 0.00316 but improved base-pair F1 from 0.476 to 0.635. The MFE and centroid methods returned the same structure for this sequence. The Mountain Centroid structure was closer in mountain profile to the reference than the structure returned by both MFE and centroid (NMSMD 0.00316 versus 0.00449), whereas that structure had higher base-pair F1 (0.769 versus 0.635). The profile panels show that both the path relaxation and Mountain Centroid remain close to the reference nesting-depth pattern despite their different base-pairing patterns.

### Figure S6. Structural comparison for the 5S rRNA example

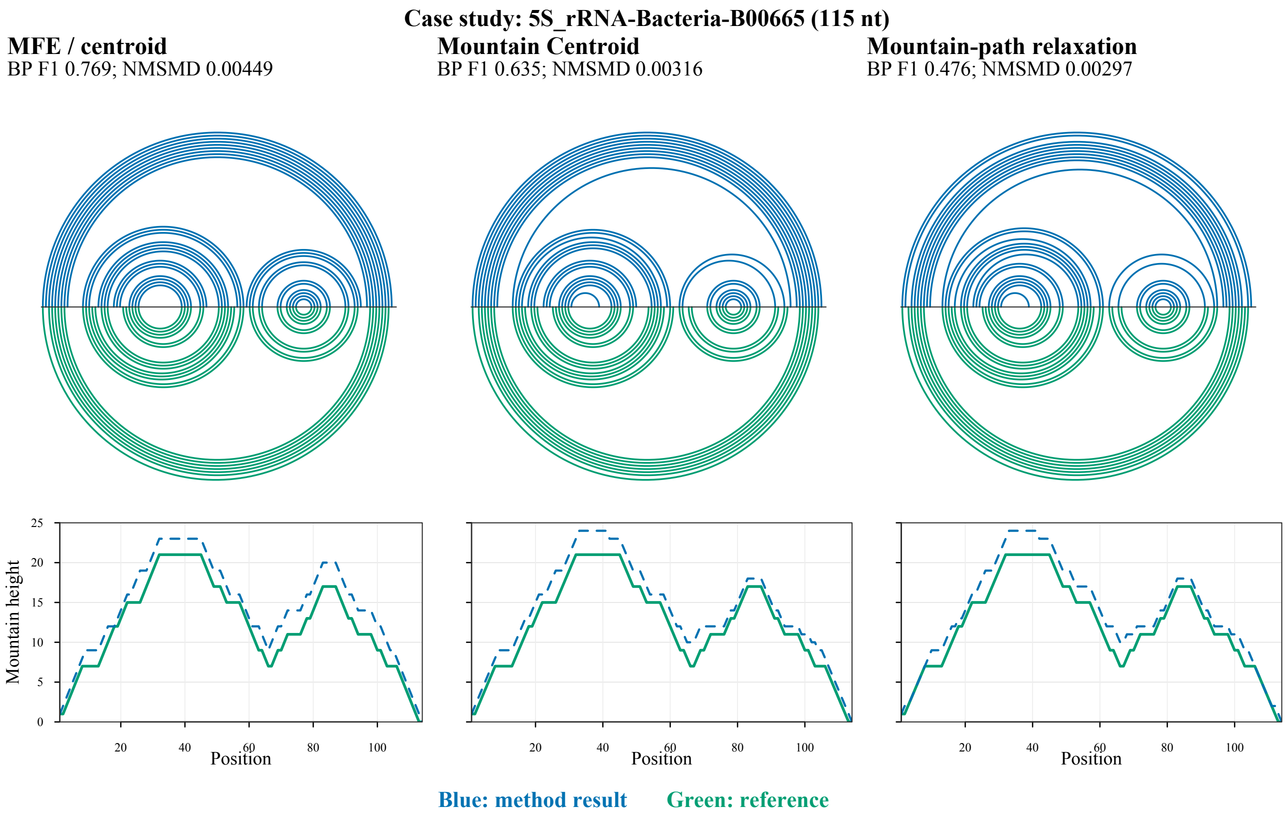


Figure S6: Structural comparison for the 5S rRNA example. Top: each arc panel shows the method result above the sequence axis and the same reference secondary structure below it. Bottom: the corresponding mountain profiles for the method result and reference. Blue denotes the method result and green denotes the reference; dashed lines distinguish method results in the profile panels. The path relaxation contained nine pairs that violated nucleotide pairability and none that violated the minimum-hairpin requirement. For this sequence, ViennaRNA MFE and centroid returned the same structure. Arcs were rendered with R4RNA; the software version is listed in Table S1.
